# Mouse xenograft model of Corticospinal Tract by Delayed Transplantation of Olfactory Ensheating Cells in Adult Rats

**DOI:** 10.1101/477521

**Authors:** Maryam Naghynajadfard, Ying Li, Geoffrey Raisman

## Abstract

Adult rats were trained to use their forepaw for retrieving a piece of noodle through a slit in the front of the cage. The dorsal corticospinal tract was lesioned by a focal stereotactic radio-frequency lesion at the level of the first/second cervical segment. Complete destruction of one side of the corticospinal tract completely prevented the use of the ipsilateral forepaw reaching for at least 6 months after operation. Rats which have shown no forepaw retrieval by 8 weeks were xenotransplanted with a suspension of cultured olfactory ensheathing cells derived from the mouse olfactory bulb, into the lesion site. Starting between 1 and 3 weeks, 10 rats with transplants bridging the lesion site resumed ipsilateral forepaw reaching. The histology of the lesioned rats with misplaced olfactory ensheathing cell showed no functional recovery during the 8 weeks of training.

## Introduction

It has been reported previously by Jike. Lu, et al., (2002) “delayed allograft transplant of Olfactory Ensheathing Cells (OEC) in rat unilateral lesion of the dorsal corticospinal tract (CST) mediate directed forepaw retrieval by ipsilateral forepaw″. Base on Ying Li et al., (2003) “The potential value of such an approach to human spinal cord injuries is the repair of delayed cervical spinal cord by 3 months″. However as recent studies show the use of auto and allo-transplant of OECs is not practicable for therapeutic use due to inaccessibility to olfactory bulb and unavailability of donors. Also although some scientists use periphery derived OECs, Olfactory Mucosa Cell (OMCs), obtained from small biopsy of olfactory mucosa lamina propria from external nares (nostrils), it is not efficient alternative source is not feasible as the auto and allo-transplant of OM in solid piece and cultured OM cells does not have enough cell numbers (Jike Lu, et al. 2002). At this study, we look at the xenotransplant of olfactory ensheathing cells derived mice olfactory bulb as the OEC cell source in the rat unilateral corticospinal tract lesioned model to retrieve Directed Forepaw Reaching (DFR) with the hope of future therapeutic use of xenotansplant source of OEC cells as an alternative stem cell source to solve some problems such as inaccessibility and unavailability of cell numbers of auto-transplantation of OEC and OM and also the difficulty to find the donor for allograft-transplantation.

## Materials and Methods

*Cell cultur.* The instruments were autoclaved by and left in flow hood. Growing media was made by (50:50) DMEM/F12 (Invitrogen, Uk) media with penicillin p/s and 10 % foetal calf serum (FCS) and left with few dishes of hank in the cold plate in flow hood. Animal were placed under terminal anaesthesia and decapitate by CO2 chamber. According to Jun Wu. et al., 2008 “The olfactory bulb was dissected out and placed in a 60 mm dish containing hanks (HBSS) (life Technologies, Uk)″. Under dissecting microscope as much of the meninges and cartilage were removed as possible using no 5 tweezers. The glomeruli were dissected out from the olfactory bulbs by slicing the bulbs in half down the centre. As Sinead M. G., et al, 2009 explained “Using small blue curved tweezers and a scalpel, white matter was removed and the tissue was washed gently in hanks and transferred into 15 ml tube with 2ml of hanks with 200 μl of 1% of 25%Trypsin-EDTA (Worthington, UK) and left in oven at (37°C, 5% CO2) for 15 mins”. After trypsinization, 10 ml of growing medium was added to stop trypsinization process. After the tissue was settled, the media was removed and only 1.5 ml media in bottom and 2 μl of DNAse (Invitrogen, Uk) solution was added and triturated by pipette tip until no big lumps remained. More media was added and if there were any lumps at bottom, cell suspension transferred to fresh tube spined at 1200 rpm for 5 mins, supernant removed and 2 ml of medium was re-suspended and plated them down on PdL coated dishes. Cultures are fed and medium is changed after 4-5 days and thereafter every 2-3 days for 16-18 days.

## Result

The cross and horizontal sections of GFAP and NF staining of 10 animals which have the DFR function return shows the regenerated CST fibres in the lesion area after 8 weeks postoperative. Figure 5 shows the elongated NF stained axons on the right hand side of the lesion however the CST fibers has not been traced to look at the terminal points in Gray matter. The figure 4 shows the cross section of the GFAP labeled astrocytes and NF stained axons in the lesion area which is almost 1% of total axons. Also The histology images show that the animals after two weeks start immunorjecting the xenotransplanted mice OECs in grafted tissue. Among five groups of animals the one who had survival time of 4 and 1 week did not show any immunorejection and the GFP labelled OEC cells can be visualised and there is not inflammation sign. However the histology images of groups 3, 4 and 5 animals with the survival time of 10-14 days,3, 4 and 5 weeks show immune reaction to xenogenic transplant of OEC degenerated cells and the central necrosis and lymphocyte infiltration in lesion area (Fig 3) . Moreover, the images of postoperative xenotransplanted OECs under daily subcutaneous injection of 100 ml of a dosage of 6 mg/kg/day avoid immunorejection e.g (fig4) which shows the misplace of GFP labeled transplanted OECs with the survival time of 4 weeks and there is no sign of inflammation.

**Figure 1:**
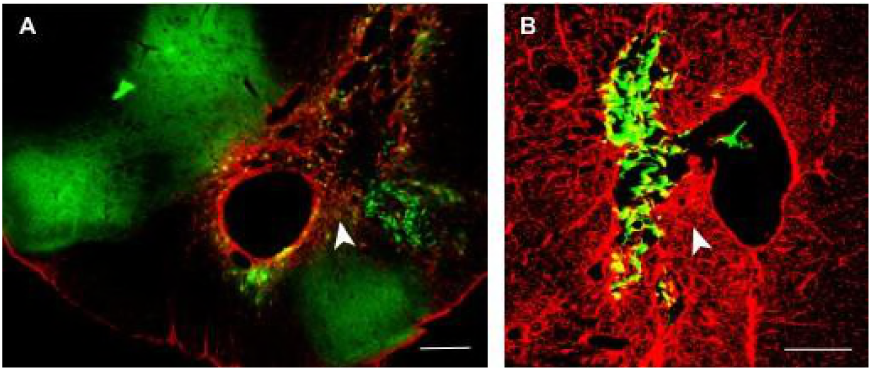
The 20 μm thickness coronal section of transplanted CST lesion failure. A. The image shows the GFP labelled grafted OECs at the dorsolateral side of lesion. B. The enlarged view of transplanted side show the distance between OEC transplanted and the lesion. Surviving time: 4 weeks; scale bar: A, 100 μm; B, 50 μm.

**Figure 2:**
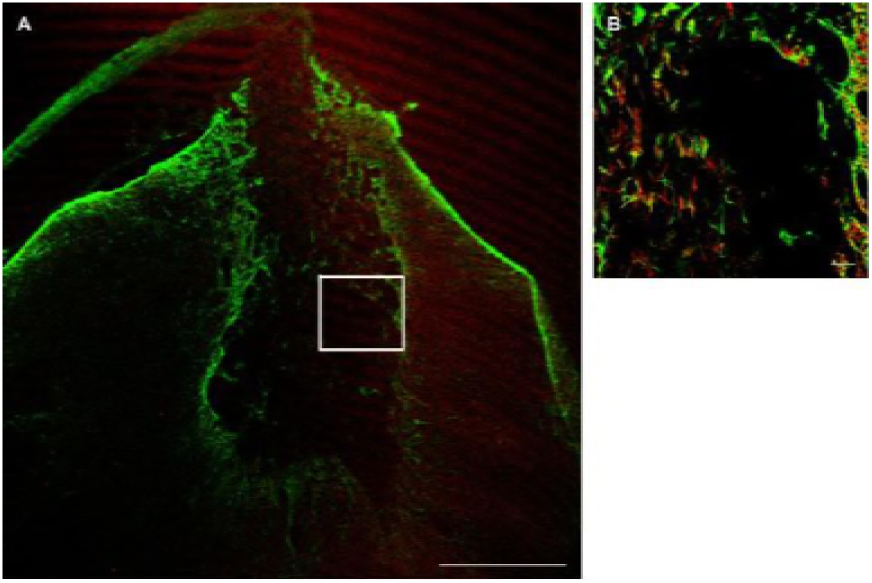
A. The regenerated Axon after transplant of olfactory ensheathing cells.B. The image of astrocytes immunostained with an antibody against glial fibrillary acidic protein (GFAP) and axons labeled with red labelled neurofilament antibody, 20μm-thickness. Survival time: 4 months, Scale bar: A; 150μm B; 60μm

**Figure 3:**
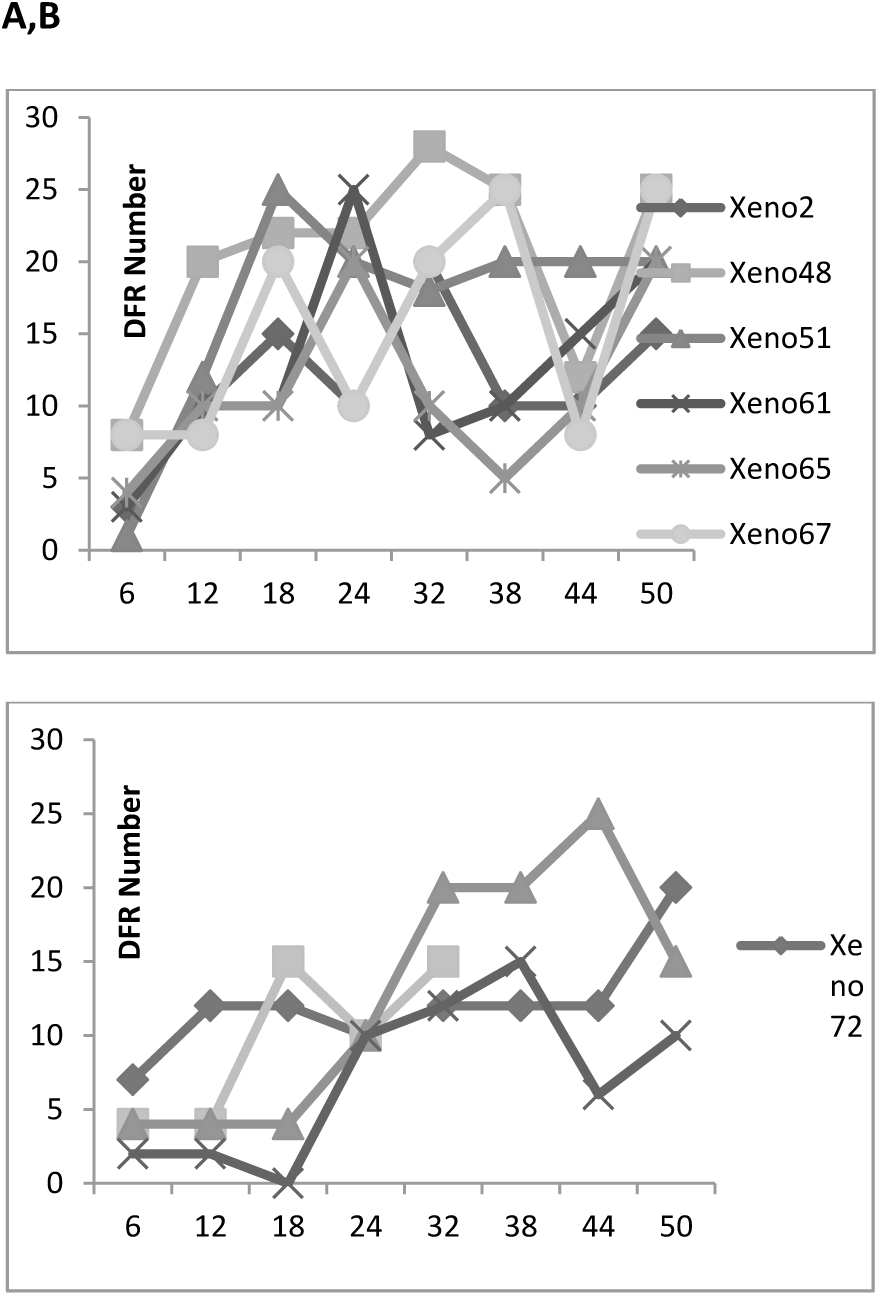
A and B Changes with time in weeks (*x*-axis) in the number of retriel (y-axis) performed by the forepaw ipsilateral to the lesion in each testing period (of a total of 50 retrievals by both forepaw.

**Figure 4:**
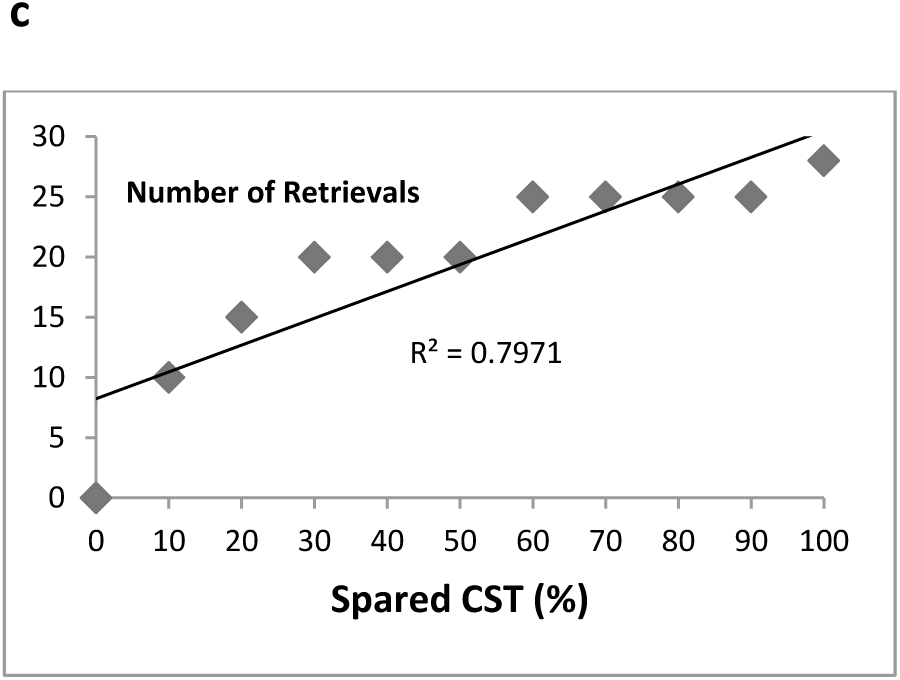
A and B Changes with time in weeks (*x*-axis) in the number of retrievals (y-axis) performed by the forepaw ipsilateral to the lesion in each testing period (of a total of 50 retrievals by both forepaws). The rate of recovery of DFR over the 8 postoperative weeks is proportional to the amount of the CST spared by the lesion. B. Significant correlation (R^2^ = 0.79) between the total number of retrievals by the ipsilateral forepaw (y-axis) and the proportion of the CST spared (x-axis).

## Discussion

### The effect of CST Lesion on DFR function

The total number of 93 animals after being trained for DFR task were under unilateral CST lesion operation using 0.25 diameter built-in thermocouple (TC) temperature sensor electrode. In total 20 rats lost the ability to retrieve their paw unilaterally. As the behaviour test results show these animals lost the ability to reach their paw through the cage slot to grasp the noodle when the number of reaching were counted 30 times three times a week over the 8 weeks postoperative. Therefore animals show 0 times retrieval. This result correlates with histology analysis of lesioned tissue. As you can see at(Figure 2,B&C) the lesion we produced at the CST tract unilaterally destroyed dorsal CST fibers and slight damage to ascending (sensory) dorsal columns totally damaged ipsilateral DFR function and no function return for the maximum period of 12 months. According to the study of Mckenn. et al., 2000 on proteoglycan effect on repair of laminin–mediated axon growth, the loss of DFR is associated with impairment of rostral motor neuron innervating proximal muscle which leads to failure in the ability of proximal muscular to extend the forelimb through the slit and it would not cause any failure in grooming, handling food, grasping and walking of the affected forelimb. According to the observation we made at the present study, the lesion we produced at the CST tract unilaterally destroyed dorsal CST fibers and slight damage to ascending (sensory) dorsal columns totally damaged ipsilateral DFR function and no function return for the maximum period of 12 months.

### Restoration of function by delayed transplantation

In future clinical trial of spinal cord injury xenograft transplant of OECs will be more efficient than allogenic transplant of human OECs from patient olfactory bulb is not feasible and the autotransplant or allotransplant of olfactory mucosa cells is not accessible due to difficulty of finding donors and the problem with the cell number of cultured tissue. At the present study we design rat delayed xenograft transplant model to investigate regeneration of CST fibres and DFR function return after transplanting cells from mice bulb and the effect of stopping immunosuppressant agent (cyclosporine) in DFR function . Recovery of function accrued if the transplanted OECs injected unilaterally in the CST lesion area and the immune attack being suppressed by injecting immusuppressant reagent regularly. As it has been explained in chapter 3 after OEC xenotransplant of total 39 AS rats, 29 rats did not show any DFR function recovery of the lesioned paw due to either misplace of cell transplant in CST dorsal column e.g (fig. 4) the cells being injected 1-2 mm away from the dorsolateral side of the lesion or the immunorejection of CST cells which can be due to the error in the injection of immunosuppressant. Of those xenotransplanted rats only 10 rats have recovered from DFR deficit and being able to reach their lesioned paw through the aperture to grasp the a noodle and their DFR ability continued rising up to 8 weeks after transplantation. As the immunogistochemistry analysis proved the xenotransplanted OEC cells migrate caudally in to distal CST and by 10 days postsurgery they form ~100-150 μm elongated bridge alongside the nerve axis and allows the CST fibres to cross the lesion area and terminated in dorsal horn e.g (fig1 A,B) . As the results show although we stop daily cyclosporine injection by the end of 3 weeks, the DFR function remains constant as it was prior stopping cyclosporine however; no GFP labelled OEC was visualised under florescent microscope due to immunorejection e.g.(fig2 A&B). This immunohistochemistry results correlate with numerical data of behaviour test as you can see in (fig3,A&B) the number of DFR retrieval after transplanting OEC starts increasing and reached to maximum number of 20 -30 during the first 3 weeks post operative under cyclosporine injection and remains spontaneous up to 4 weeks after stopping cyclosporine injection and although the histology image proved that by stopping cyclosporine xenotransplanted OECs being rejected at week 8 of posts surgery, the immunorejection of grafted tissue did not effect on DFR Recovery and numerical data shows that the animals did not shows deficit only in their paw reaching task and the number of DFR remains spontaneous between 10-30 . This result is also consistent with the study of David, Choi. et. al., (2003) who studied on facial nerve xenograft model. The histology analysis shows that ~ 1% of the total 50,000 myelinated CST fibres on OEC cells transplanted animals (fig .3 B) shows that regenerated fibres are ~ 1% of the total 50,000 myelinated CST fibres on OEC cells transplanted animal and this result is correlated with the number of spared CST fibres with R^2^ = 0.79 (fig 4). These results are also consistent with previous studies by Li et al., 1997 and Naghmeh N., et al., 2003 in allogenic transplant of OEC in unilateral lesioned CST model, our observation in cross-species transplantation immunorejection indicates that transplanted rats start immurejecting OEC cells by 10 days postoperatively. The histological evaluation by haematoxylin-eosin of graft biopsy specimens show tissue necrosis and inflammatory cell infiltration in lesion area which according to the Banff classification of inflammatory reaction was accepted as grade III. However, clinical evaluation of transplanted animal did not show any signs of rejection such as Erythema, odema and hair loss (Cendales LC. et al., 2008).

## Conflict of Interest

Corresponding Author Declaration Form Manuscript title Mouse xenograft model of Corticospinal Tract by Delayed Transplantation of Olfactory Ensheating Cells in Adult Rats Corresponding author: Maryam Naghynajadfard Additional authors in the order provided in the manuscript: Ying Li The corresponding author must provide statements of authorship, originality, conflicts of interest, and research funding on behalf of all authors of the manuscript. Corresponding author, on behalf of all authors, is equired to report the following information with each submission: 1. All third-party financial support for the work in the submitted manuscript. 2. All financial relationships with any entities that could be viewed as relevant to the general area of the submitted manuscript. 3. All sources of revenue with relevance to the submitted work who made payments to you, or to your institution on your behalf, in the 36 months prior to submission. 4. Any other interactions with the sponsor of outside of the submitted work should also be reported. 5. Any relevant patents or copyrights (planned, pending, or issued). 6. Any other relationships or affiliations that may be perceived by readers to have influenced, or give the appearance of potentially influencing, what you wrote in the submitted wo rk. As a general guideline, it is usually better to disclose a relationship than not. This information will be acknowledged at publication in a Transparency Document. Additional information on the ICMJE recommendations can be found at: http://www.icmje.org. The corresponding author must complete the Corresponding Author declaration form on behalf of all authors of the manuscript.

The corresponding author signed this statement on behalf of all co-authors to indicate that the above information is true, correct and complete. Signature: Naghynajadfard Print name: Maryam Naghynajadfard Date: 11/22/2018

